# Reduced neural sensitivity to musical tempo despite enhanced neural tracking of musical features in older adults

**DOI:** 10.1101/2025.03.24.644846

**Authors:** Yue Ren, Kristin Weineck, Molly J. Henry, Björn Herrmann

## Abstract

A substantial body of prior research has focused on how aging affects the neural processing of speech, whereas less is known about how older adults encode naturalistic music. Here, we investigated whether the neural tracking of different features in naturalistic music differs between age groups. Younger and older adults (both sexes) listened to excerpts of naturalistic music with different tempi (1-4 Hz) while electroencephalography (EEG) was recorded. The results show an age-related enhancement of neural responses to sound onsets, suggesting a loss of inhibition in the aged auditory cortex that is thought to arise from peripheral decline. This hyperresponsiveness generalized to the neural tracking of multiple features (amplitude envelope, onsets, beat, and spectral flux) in naturalistic music. Crucially, older adults showed reduced sensitivity of early brain responses (0-130 ms) to tempo: Unlike younger adults, whose neural tracking decreased systematically with increasing tempo, older listeners maintained uniformly enhanced tracking across tempi. Spectral flux best predicted tempo- related changes in EEG activity, but this was reduced in older compared to younger adults. In sum, the current study demonstrates that cortical hyperactivity in aging enhances the tracking of different features during naturalistic music listening but impairs the sensitivity to musical tempo. This dissociation highlights the complex changes and functional consequences of auditory aging related to the processing of naturalistic music.

**SIGNIFICANCE STATEMENT:** How aging affects the encoding of ecologically-valid, naturalistic music is not well understood, although music is an important part of older people’s lives. Here, we used electroencephalography and temporal response function approaches to investigate how younger and older adults encode different features of naturalistic music at different tempi (1 to 4 Hz). The results reveal a fundamental trade-off in the aging auditory system: enhanced neural tracking of acoustic features coexists with declined sensitivity to musical tempo. This dissociation between hyperactivity for feature encoding and impaired temporal sensitivity highlights complex changes in the aged auditory system that can have implications for music perception in older adulthood.

## INTRODUCTION

Aging is associated with a host of changes in auditory function, including reduced sensitivity to sound due to peripheral decline (Gratton and Vázquez, 2003; Jayakody et al., 2018) and aberrant plasticity in brain regions downstream, most prominently in auditory cortex (Ibrahim and Llano, 2019; Herrmann and Butler, 2021). Age-related changes in auditory function affect the processing of complex sound stimuli, with a large body of work focusing on changes in the perception and neural encoding of speech (Gordon-Salant and Fitzgibbons, 1999; Wöstmann et al., 2015). However, age-related changes in auditory function likely also impact music processing. For instance, older adults show different music preferences relative to their younger selves (Cohrdes et al., 2017), exhibit declines in pitch discrimination and chord categorization (Clinard et al., 2010; Bones and Plack, 2015), and experience challenges in processing mistuned harmonics of complex tones (Alain et al., 2012). Fundamentally, tracking dynamic temporal and spectral changes in music is vital for music perception (Stewart et al., 2006; Koelsch, 2011; Janata, 2015), but little is known about how aging affects the brain’s ability to track features of naturalistic music over time.

Despite significant advances in understanding neural responses to music (Doelling and Poeppel, 2015; Norman-Haignere et al., 2015, 2022; Zuk et al., 2021), gaps remain in how the brain processes complex, naturalistic musical stimuli. Much research has focused on the neural processing of simplified music, such as tones or MIDI sound sequences. Analyses have typically relied on auditory evoked potentials (ERPs) to characterize fundamental processing mechanisms, including temporal coding fidelity via P1-N1-P2 components (Koelsch et al. 2000; Shahin et al. 2003) and deviance detection (encompassing spectral, temporal, or intensity changes) through the mismatch negativity (Näätänen et al., 1987; Tervaniemi et al., 2001). Despite the advantages in isolating specific acoustic features and establishing a relationship between stimulus properties and neural responses (Näätänen et al., 2007), these stimuli and ERP approaches are unable to capture the complexities and temporal evolution of real music.

Research addressing this ecological gap has focused on neural tracking, that is, the synchronization of neural activity (e.g., measured via EEG/MEG) with dynamic features of continuous stimuli (Lalor and Foxe, 2009; Lalor et al., 2009). Recent work has focused on the neural tracking of the amplitude envelope in naturalistic music (Di Liberto et al., 2020; Zuk et al., 2021; Keitel et al., 2025), inspired by speech-envelope tracking research (Luo and Poeppel, 2007; Ding and Simon, 2012). However, spectral features (e.g., timbre and melodic structure) are equally crucial in music. Research demonstrates that younger listeners track spectral content more robustly than the amplitude envelope of naturalistic music, with strongest tracking at slower tempi (1-2 Hz; Bauer et al., 2015; Weineck et al., 2022). This tempo-effect may be linked to auditory event density, where sparser temporal structure allows more effective neural encoding (Näätänen et al., 1987). This research has thus far only focused on younger adults, whereas the extent to which aging affects the neural tracking of these features during naturalistic music listening is unknown.

Aging and hearing loss can lead to brain plasticity that may impact the neural encoding of music. Specifically, any peripheral damage that reduces acoustic inputs can lead to a loss of inhibition and an increase in excitation in the auditory cortex (Parry et al., 2018; Salvi et al., 2016; Herrmann and Butler, 2021). This, in turn, can manifest as hyperresponsiveness to sound, where, for example, simple tones or noise bursts often elicit larger ERP responses in the auditory cortex of older people compared to younger adults (Alain et al., 2012; Presacco et al., 2016). Moreover, older adults show increased neural tracking of amplitude and spectral features in simple continuous stimuli (e.g., tone sequences, modulated noise; Purcell et al., 2004; Goossens et al., 2016, 2019; Herrmann et al., 2019; but see Henry et al., 2017), suggesting that the loss of inhibition could also alter the neural tracking of features in naturalistic music.

A few studies have focused on the age-related changes in neural tracking of naturalistic auditory stimuli, namely the tracking of speech. The neural tracking of the amplitude envelope of speech is enhanced in older compared to younger adults (Presacco et al., 2016; Brodbeck et al., 2018; Decruy et al., 2019, 2020; Broderick et al., 2021; Panela et al., 2024), consistent with a loss of inhibition and an increase in excitation in the auditory cortex of older adults (Herrmann and Butler, 2021). However, hyperactivity may also impair the accurate encoding of features in music, which could underlie changes in music perception in older adulthood. In the current electroencephalography study, we investigate whether aging affects the neural tracking of different features (i.e., amplitude, spectral, beat) in naturalistic music at different tempi. Our multi-feature approach affords answering whether age-related hyperactivity is present during music encoding and whether it uniformly alters music tracking across different features and tempi.

## MATERIALS AND METHODS

### Participants

32 younger adults (22 females, 10 males; mean age: 24.4 years, age range: 19-34 years; data from our previous study; (Weineck et al., 2022)) and 34 older adults (16 females, 18 males, mean 66.2 years, 58-82 years of age) with self-reported normal hearing participated in the study. We reanalyzed the data from younger adults (Weineck et al., 2022) jointly with the new data from older adults that were recorded using the same experimental procedures. Data from 5 additional younger adults and 4 additional older adults were recorded but not included in further analyses, because a high number of trials contained artifacts in the EEG data (see below). Most participants self-reported normal hearing (4 participants reported occasional ringing in on or both ears and one reported hyperacusis since a young age). Participants’ demographic background was obtained online through the LimeSurvey platform (LimeSurvey GmbH, Germany, https://www.limesurvey.org) prior to the main in-person session. Participants were remunerated for their online and in-person participation (Online: 2.50 €, EEG: 7 € per 30 min). All participants provided informed consent prior to the experiment. The study was approved by the Ethics Council of the Max Planck Society Ethics Council in compliance with the Declaration of Helsinki.

### Pure-tone audiometry and estimation of sensation level

Participants underwent an audiogram hearing test (MA 25 Audiometer, Robert Bosch GmbH, Germany) at the beginning of the experiment. Audiogram data from younger participants were not available in our previous study (Weineck et al., 2022). Therefore, in the current study, we collected audiogram data from 18 new younger participants matched for age with the younger group in the original study from which the EEG data were taken (Weineck et al., 2022). Audiometric thresholds were assessed for pure tones at frequencies of 0.25, 0.5, 1, 2, 4, and 8 kHz (Figure 1A). The pure-tone average (PTA) was calculated as the average across the thresholds of 4 frequencies (0.5, 1, 2, and 4 kHz; Figure 1B). Compared to younger individuals, older adults showed elevated hearing thresholds (t_50_ = 6.16, p = 1.23·10^-7^, η^2^ = 1.80), aligning with previous research (Peelle et al., 2011; Slade et al., 2020) and the prevalence of hearing thresholds in community-dwelling older adults (Cruickshanks et al., 1998; Feder et al., 2015; Goman and Lin, 2016).

**Figure 1.**
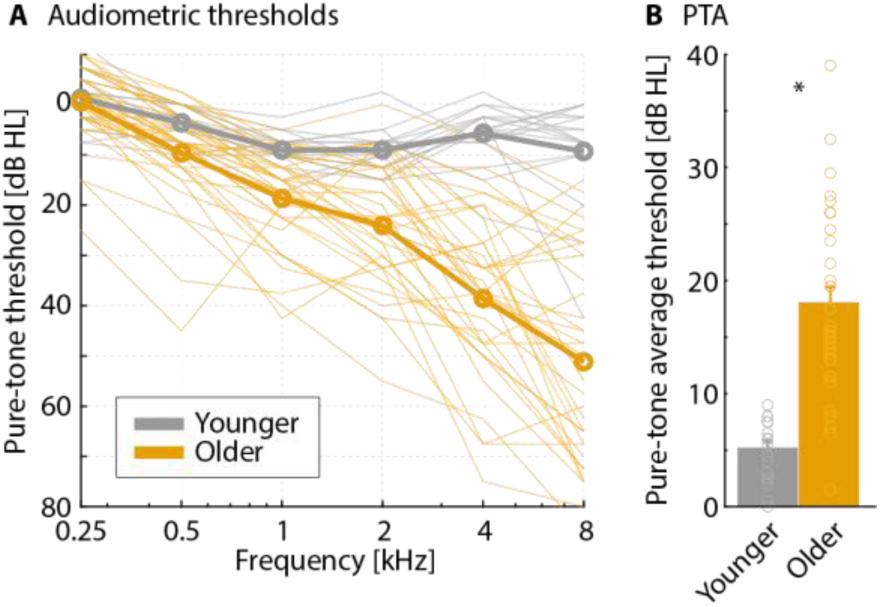
Audiograms and hearing thresholds for younger and older adults. A: Audiograms. Thick lines denote group-mean data and thin lines represent individual participant data. B: Pure-tone average (PTA) for both age groups (mean across 0.5, 1, 2, and 4 kHz). Individual data points are denoted by dots. *p < .05.

The sensation level was also determined for each person in MATLAB. To this end, participants listened to white-noise sounds of 12 s duration whose sound level continuously increased or decreased (6 trials each), and responded promptly upon detecting the presence of the noise (increasing sound level) or no longer hearing the noise (decreasing sound level) (Herrmann and Johnsrude, 2018). The sound level at which a participant pressed the button was averaged across the 6 trials to obtain the participant’s sensation level. Note that sensation- level data from young adults were available from the previous study (Weineck et al., 2022). Sensation levels were lower for younger adults (-63.1 dB; MATLAB full scale) compared to older adults (-52.6 dB; t_64_ = 6.95, p = 2.31·10^-9^, η^2^ = 1.71), as expected given the audiograms.

For younger adults, all music stimuli were presented at 50 dB above their individual sensational level (Weineck et al., 2022). Music stimuli for older adults were presented at a fixed sound level that corresponded to the mean level at which sounds were presented to younger adults. Hence, sounds were, on average, presented at the same sound-pressure level (SPL) for younger and older adults. We opted to present stimuli at the same SPL rather than at the same sensation level (that is, relative to individual thresholds), because the latter would have led to higher sound intensities for older compared to younger adults, given their elevated thresholds. Higher sound intensities can lead to larger brain responses (Woods et al., 2006; Sun et al., 2012). This could have confounded our main hypothesis that the neural tracking of music may be enhanced for older compared to younger adults, given previous age-related neural tracking enhancements for speech and modulated noise sounds (Alain, 2014; Herrmann et al., 2019). By presenting music stimuli at the same SPL across age groups (on average), we rather work against this hypothesis, while ensuring that stimuli were clearly audible for all participants. Using the same SPL also reflects more closely natural environments where the sound level is often not under control of a person. Moreover, older adults may have a compressed dynamic range, such that very soft sounds are not perceived, whereas sound at high levels can be perceived as being too loud (Slade et al., 2020; Shehabi et al., 2022). Younger and older adults tend to perceive sounds as similarly loud for the mid-range sound levels used in the current study (Herrmann et al., 2019).

### Stimuli

We adopted the same stimuli and procedures that we used to record the data for younger adults in our previous study (Weineck et al., 2022). The auditory materials initially consisted of 93 segments extracted from 39 full instrumental musical pieces spanning four primary genres: techno, rock, blues, and hip-hop, accessible for download from the Qobuz Downloadstore (https://www.qobuz.com/de-de/shop). Each segment was manually epoched at musical phrase boundaries, such as transitions between choruses and verses, resulting in musical segments with durations varying from 14.4 to 38 s. To determine the original tempo of each musical segment, we did not use beat counts from publicly available software, as its outputs lacked sufficient reliability for cross-genre tempo analysis. Instead, we asked a professionally trained percussionist to tap to the beat of each segment with a wooden drumstick on a MIDI drum pad. The tap tempo (BPMs, beats per minute) was extracted from the MIDI files. Each segment was repeated three times, and the final BPM was the average between the two takes with least difference.

Musical segments were then tempo-modulated to a range of 1 to 4 Hz in steps in 0.25 Hz increments (i.e., 13 tempi). The tempo manipulation was carried out using a customized MAX patch (MAX 8.1.0, Cycling ’74, CA, USA), which changes the tempo of music without changing the pitch. To ensure that stimuli had unambiguous, easily tappable beats, 8 individuals were asked to listen and tap to each segment on their computer keyboards for a minimum of 17 taps while their key strokes were recorded and analyzed using an online BPM tool (beats per minute; https://www.all8.com/tools/bpm.htm). 21 musical excerpts were excluded exhibiting high inter-tapper variability (>2 tappers diverging from the group) or poor sound quality. The process resulted in 703 music excerpts, varying in duration from 8.3 to 56.6 seconds across the 13 tempi conditions. While faster tempi resulted in shorter durations and slower tempi in longer durations for a unique musical piece, this ensures that the same number of rhythmic cycles are present for each condition.

To obtain a balanced distribution of stimulus conditions and avoid redundancy, the musical excerpts were pseudo-randomly allocated into four groups (Kaneshiro et al., 2020). Each group comprised 159 to 162 musical excerpts (trials). Each musical excerpt could repeat up to three times across different tempi within the same group, but was never repeated for a unique tempo (Weineck et al., 2022). All musical excerpts were mono WAV files with a sampling rate of 44,100 Hz. Each file featured a smooth linear-ramping fade-in and fade-out phase of 500 ms, and underwent root-mean-squared normalization using custom MATLAB scripts. Auditory stimuli in the study were played through a Fireface sound card (RME Fireface UCX Audiointerface, Germany) via headphones (Beyerdynamics DT-770 Pro, Beyerdynamic GmbH & Co., Germany). Auditory and visual stimuli were delivered using custom-written MATLAB scripts with Psychtoolbox (PTB-3, Brainard, 1997) in MATLAB (R2021a; The MathWorks, USA).

### Experimental design

The experimental procedures were conducted in an acoustically and electrically shielded booth. Verbal instructions for the experimental procedures were provided by the experimenter, complemented by written instructions displayed on the screen (BenQ Monitor XL2420Z, 144 Hz, 24”, 1920 × 1080). Each participant was assigned to one of the four groups of musical excerpts, with total trial numbers varying from 159 to 162. The study lasted approximately 2.5 hours.

The main task comprised eight blocks, and participants were allowed to take breaks between blocks as needed. Each block contained 19 to 21 trials, depending on the group of stimuli to which the participant assigned. Each trial comprised three phases: attentive listening (music stimulation without movement or responses), tapping (music stimulation with finger tapping) and rating. In the attentive listening phase, a fixation cross was presented on the computer screen and a music excerpt was presented to the participant. Participants were instructed to mentally identify the beat of the musical excerpt without any physical movement, devoting full attention to the music presentation. The duration of this phase varied for each trial, ranging from 8.3 to 56.6 seconds (depending on the duration of the musical excerpt). Following a 1- second interval after the music presentation ended, a visual cue (a cartoon picture of a hand) appeared on the screen, signaling the start of the tapping task. Simultaneously, the final 5.5 s segment of the previously heard musical excerpt was replayed, and participants were instructed to tap a finger in accordance with the beat. In the rating phase, participants were asked to assess the musical excerpt separately based on enjoyment/pleasure, familiarity, and ease of tapping to the beat using a visual analogue scale with the computer mouse. The scale ranged from -100 to +100, representing self-evaluations from ‘not at all’ to ‘very much’. Participants performed a brief three-trial training before the main task was performed. The aim of the current study was to investigate the degree to which the neural encoding of different features and tempi in naturalistic music differed between age groups. Analyses thus focused on the ‘attentive listening’ phase (i.e., phase 1) of the experiment.

### EEG recording and pre-processing

EEG data were recorded using the BrainVision Recorder (v.1.21.0303, Brain Products GmbH, Gilching, Germany) and a Brain Products actiCAP system with 32 active electrodes attached to an elastic cap based on the international 10–20 placement (actiCAP 64Ch Standard-2 Layout Ch1-32, Brain Products GmbH, Germany). The EEG signals were sampled at a rate of 1000 Hz using an online low-pass filter at 250 Hz with electrode impedances consistently maintained below 10 kOhm. To ensure precise time alignment between the recorded EEG data and the auditory stimuli, a TTL trigger pulse was sent at both the onset and offset of each musical excerpt via a parallel port.

Offline data analysis was carried out using MATLAB software (R2021b; The MathWorks, Natick, MA, USA) with custom-written code combined with the Fieldtrip toolbox (Oostenveld et al., 2010). The continuous EEG data were bandpass filtered between 0.5 and 30 Hz (Butterworth and FIR two-pass filter), re-referenced to the average reference, and down- sampled to 500 Hz. EEG data were segmented into time series time-locked to the onset of each musical excerpt. Trials exhibiting a signal change of more than 1000 μV for any electrode were excluded from analyses to remove trials with huge artifacts that could bias Independent Component Analysis (ICA). ICA was calculated to identify and suppress activity related to eye blinks, horizontal eye movements, and cardiac artifacts. Following the transformation of data back from the ICA component space to the electrode space (after removing artifact components), trials with a signal change exceeding 400 μV in any electrode were excluded from analyses. Participants with less than 60% of valid trials were removed from analysis. Data from 5 younger adults and 4 older adults were excluded based on this criterion.

### Data analysis

#### Analysis of neural response to music onset

We first examined the neural responses elicited by the onsets of the music stimuli. This analysis was conducted to investigate whether hyperactivity is present in older compared to younger adults (Anari et al., 1999; Herrmann and Butler, 2021). For each participant, EEG data were divided into epochs ranging from 100 ms before to 400 ms after the onset of each musical excerpt. Data were averaged across all trials. The averaged response was baseline-corrected by subtracting the mean amplitude in the pre-stimulus time window (-100 to 0 ms) from the amplitude at each time point. Data were averaged across 10 electrodes from a frontal-central cluster (F3, F4, Fz, FC5, FC6, FC1, FC2, C3, Cz, C4) that is known to be most sensitive to neural activity originating from auditory cortex (Näätänen et al., 1987; Picton et al., 2003). To examine the difference in onset responses between age groups, we conducted independent samples t-tests on the response amplitude at each time point. We identified the time intervals indicative of group differences based on the adjusted p values (*p<.05) using the False Discovery Rate (FDR) method (Benjamini and Hochberg, 1995; Genovese et al., 2002).

#### Calculation of feature vectors in naturalistic music

We used the features adopted in our previous publication for all musical stimuli (Weineck et al., 2022). In total, four features were calculated: amplitude envelope, amplitude-onset envelope, beat, and spectral flux. The amplitude envelope was determined using a 128- channel gammatone filterbank between 60 to 6000 Hz (Patterson et al., 1987; Lopez-Poveda and Meddis, 2001). To calculate the amplitude-onset envelope, we applied half-wave rectification to the first derivative of the amplitude envelope, capturing the onsets and energy changes in the stimulus over time (Fiedler et al., 2017; Panela et al., 2024). The beat of each music segment was determined by a professional percussionist drumming in synchronization with the original tempo using drumsticks resulting in a binary-coded time series (0 for no beat and 1 for beat). The percussionist repeated each segment three times. The final beat vectors were determined by the average two takes with the minimal differences. Spectral flux was calculated by comparing the logarithmic amplitude spectrogram between consecutive time frames (each frame comprising 344 samples), reflecting the rate of power spectrum changes. All features in the naturalistic music were z-scored and down-sampled to 128 Hz. To ensure consistent sample sizes across stimulus features, the data were trimmed to match.

#### Analysis of neural tracking of features in music

We used temporal responses functions (TRFs) and EEG prediction accuracy to investigate whether the neural tracking of features in music differed between age groups. The temporal response function (TRF) is a system identification model that relates features of a sensory stimulus and the neural response based on regularized linear regression (Ding and Simon, 2012; Haufe et al., 2014). In this study, we adopted the TRF model to characterize how different features map onto the EEG signals (forward model) using the MATLAB toolbox “The multivariate Temporal Response Function (mTRF) Toolbox” (Crosse et al., 2016, 2021). To eliminate onset responses elicited by the start of the music excerpt, data from the first second following stimulus onset were excluded from further analysis. EEG data were down-sampled to 128 Hz for the TRF calculation.

For each of the 13 tempo conditions, EEG data from all trials and the corresponding time series for each feature were selected for model calculation using leave-one-trial-out cross-validation iterations. For each iteration, the EEG data and feature time series of one trial served as the test dataset and was held out, whereas the EEG data and feature time series of the remaining trials served as the training dataset. The model training incorporated time lags ranging from 0 - 400 ms (using the functions *mTRFcrossval* and *mTRFtrain*). The model implemented ridge regularization to prevent overfitting, and the training process allowed us to determine the optimized ridge parameter (λ), maximizing the correlation between the actual and predicted neural response from the training outputs (λ range: 10^-6^ - 10^6^). This process was repeated for each feature (Di Liberto et al., 2020). The primary output of the model training – that is, TRF weights assigned to each channel – reflects the modulation of the EEG signal in response to a change in a particular feature (Crosse et al., 2016, 2021) and was used for subsequent analysis. To obtain the accuracy with which the EEG response of the test dataset (i.e., the trial that was left out) could be predicted based on the TRF model, the feature time series of the test dataset was convolved with the TRF weights, resulting in a predicted EEG time series (separately for each feature). EEG prediction accuracy was then calculated as the Pearson correlation between the predicted and the actual EEG data (EEG prediction accuracy was calculated using the function *mTRFpredict*). TRF weights and EEG prediction accuracy were calculated for each participant separately for each of the 13 tempi. TRF weights and EEG prediction accuracy were averaged across the 10 electrodes of the frontal-central cluster (F3, F4, Fz, FC5, FC6, FC1, FC2, C3, Cz, C4) that are sensitive to neural activity from the auditory cortex.

We first assessed neural responses for each age group and age-group differences in neural responses independently of musical tempi. To this end, TRF time courses (weights) were averaged across the 13 tempi, separately for each feature and participant. We examined the TRF weights within each age group by calculating a one-sample test against zero, separately for each time point. FDR-thresholding was used to account for multiple comparisons. This analysis tests whether neural responses were modulated by a specific feature in music, separately for each age group. We then compared TRF weights between the two age groups using separate independent samples t-tests for each time point (including FDR-thresholding).

TRF time courses are characterized by positive and negative deflections (peaks), similar to the traditional event-related potential (Crosse et al. 2016; Crosse et al. 2021; Luck 2014). To examine response magnitude differences between age groups, analyses focused on the peaks in the TRF time courses. We analyzed the P1-N1 (first major positive peak minus first major negative peak) and P2-N1 amplitude differences in both age groups, separated by the four different features (see similar approaches for event-related potential analyses and TRF analyses: (Tremblay et al., 2001; Wagner et al., 2017; Herrmann, 2025). The P1, N1 and P2 amplitudes were determined by the local extrema (maximum for P1 and P2 components, minimum for the N1 component) in the TRF weights within a 100-ms time window that included the corresponding component. The latencies of the P1, N1 and P2 components were thus identified by the time points corresponding to these extrema. For each participant, the P1, N1 and P2 amplitudes were determined by the local extreme values within a 30-ms time window centered at the averaged latencies of each component for each age group. Subsequently, the P1-N1 and P2-N1 amplitude differences were calculated by subtracting the amplitudes of the N1 from P1, and N1 from P2, respectively. Age-group differences in the P1-N1 and P2-N1 amplitude were assessed using independent samples t-tests for each feature.

In the previous study (Weineck et al., 2022), we found that the musical tempo affected TRF accuracy: TRF accuracy decreased as tempo increased, particularly for the spectral flux feature. Additionally, tempo also played a role on the TRF weights at different time points (ranging from 0 to 400 ms). Therefore, we conducted a nuanced analysis to investigate the ‘sensitivity’ of TRF weights to music pieces with different tempi. For each feature, we first defined a sliding time window of 62.5 ms (i.e., 9 samples) centered at each time point on the time courses of the TRF weights and averaged the weight values within each time window per participant. This sliding-window averaging procedure served to smooth data and mitigate local fluctuations at each data point prior to the musical-tempo analysis. Subsequently, we calculated a linear regression for each time point of the smoothed TRF, to linearly relate the 13 tempi to the TRF weights. The resulting slope value of this regression quantifies the degree of TRF weight change as a function of musical tempo (separately for each time point). Slope values were tested against zero using a one-sample t-test, separately for each age group (younger, older) and feature (amplitude envelope, onset envelope, beat, spectral flux), to examine whether musical tempo was associated with a change in the neural responses. The t-tests were performed for each time point within each age group and feature (FDR-thresholded). Age-group differences were examined using an independent samples t-test at each time point (FDR-thresholded).

In addition to the analysis of the TRF time courses (weights), we also analyzed the EEG prediction accuracy (i.e., TRF correlation: correlation between predicted and actual EEG) – another metric that reflects the strength of neural tracking. To examine age-group differences for EEG prediction accuracy without the tempi influences, we first averaged the prediction accuracy across 13 tempi separately for each of the four features. The averaged TRF correlations were submitted to a 2 × 4 mixed-design ANOVA, with ‘Age Group’ (younger, older) as a between-subjects factor and ‘Feature’ (envelope, amplitude-onset, beat, and spectral flux) as a within-subjects factor. Post-hoc pairwise comparisons were conducted using the Bonferroni correction. To evaluate how musical tempo affects EEG prediction accuracy, we calculated a linear regression relating musical tempo to EEG prediction accuracy, separately for each participant and feature. The slope of the linear regression was tested against zero using a one-sample t-test for each feature and age group. Age-groups were compared using an independent samples t-test.

## RESULTS

### Auditory onset responses in older and younger adults

Figure 2 shows the neural responses to the onset of the musical excerpts for both age groups. Within the time range of 134-260 ms, responses significantly differed between age groups (FDR-thresholded). The response difference in this time window is the result of an enhanced and delayed N1 component, along with a diminished and delayed P2 component, in older compared to the younger adults. These results are consistent with previous research in auditory perception and aging, which may suggest differences in the information flow in auditory regions in response to complex acoustic features as well as with hyperactivity in auditory cortex of older adults (Friedman et al., 1993; Friedman, 2013; Henry et al., 2017b; Irsik et al., 2021; Herrmann et al., 2022).

**Figure 2.**
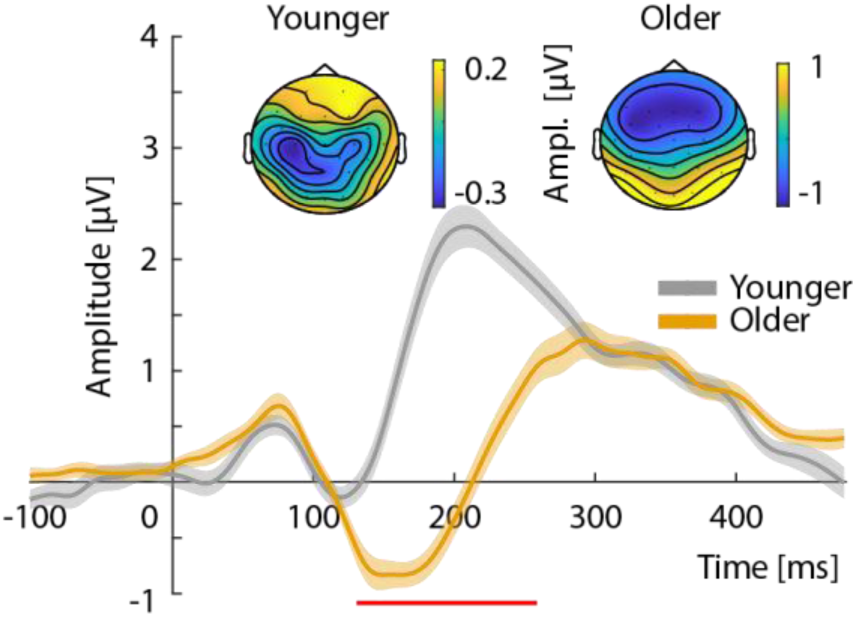
Neural responses to the onset of musical excerpts. Group-averaged time courses are displayed by the two colored curves. Shaded areas mark the standard error of the mean (SEM). The red line at the bottom marks the time window exhibiting a significant difference between age groups (pFDR < .05). Topographies for the N1 component for the two age groups (based on the time window centered on the local minimum, separately for each age group).

### Neural tracking of features in music is enhanced in older adults

We investigated the neural tracking of four different features in naturalistic music (amplitude envelope, amplitude-onset envelope, beat, and spectral flux) in younger and older adults. As shown in Figure 3, the averaged TRFs exhibited temporal patterns consistent with a canonical auditory P1-N1-P2 complex in both age groups (Martin et al., 2008; Picton, 2013). The N1 component was enlarged, and the P2 component was delayed in older adults compared to younger adults for the amplitude envelope, amplitude-onset envelope, and spectral flux, but not for the beat (p_FDR_ <.05).

**Figure 3.**
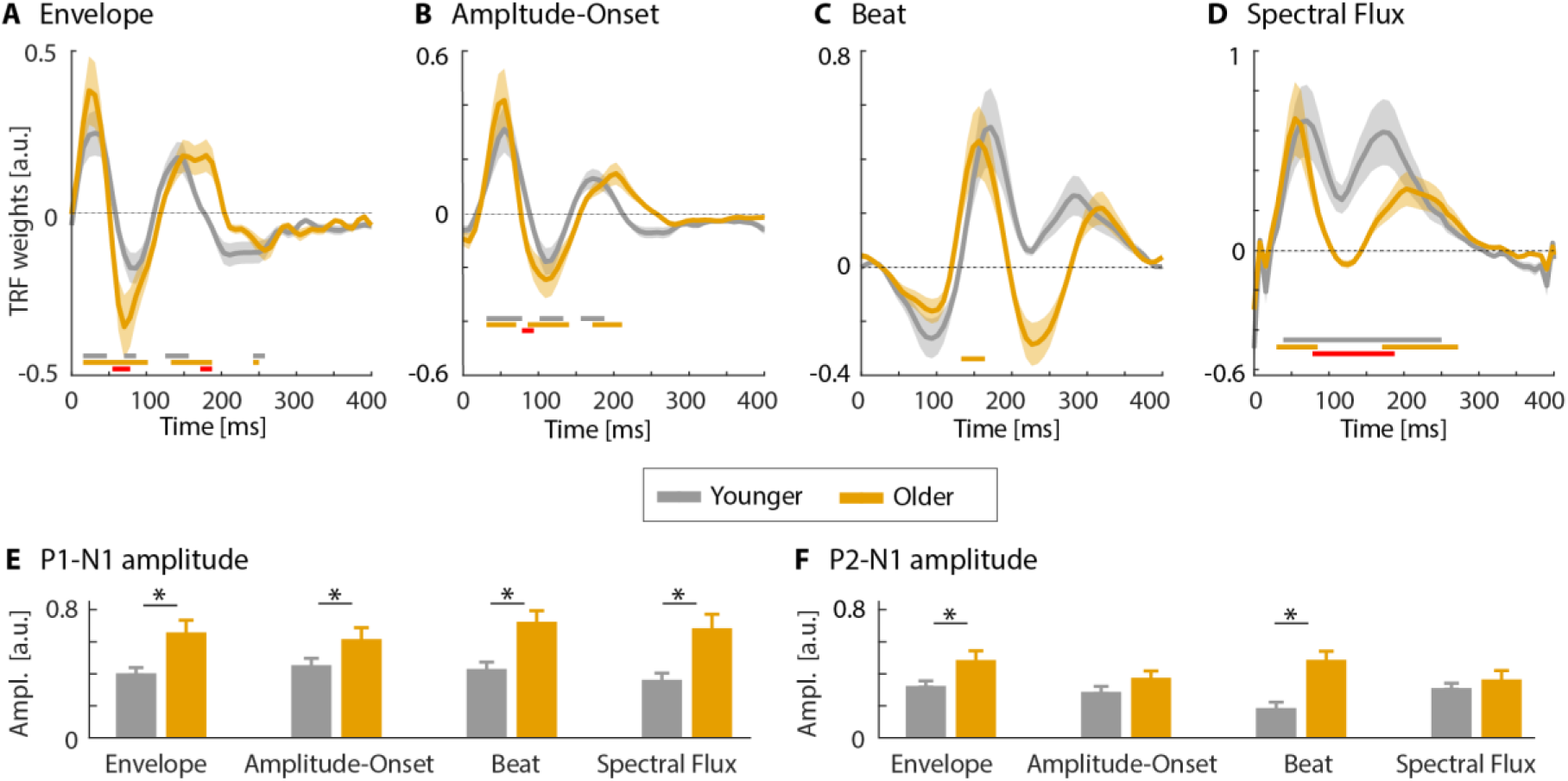
Neural tracking (TRF weights) for four features in naturalistic music and two age groups. A-D: Averaged TRF weights courses (shaded area around means reflects the SEM) for four features (averaged across tempi) and both age groups. Horizontal lines at the bottom mark the time window exhibiting a significant difference (pFDR <.05) in TRF weights between age groups (red line) and within each group against 0 (younger adults: gray; older adults: yellow). E-F: P1-N1 and P2-N1 amplitude differences for TRF weights between age groups, separately for the four features. *p < 0.05

To further investigate age-related differences in the P1-N1-P2 complex of the TRF weights, we calculated the P1-N1 and P2-N1 amplitude for each feature. Results showed that the P1- N1 amplitude was significantly larger in older adults compared to younger adults for all four features (envelope: t_64_ = 3.29, p = .002, η^2^ =.79; onset: t_64_ = 2.11, p = .04, η^2^ =.49; beat: t_64_ = 3.64, p = 5.52·10^-4^, η^2^ =.89; spectral flux: t_64_ = 3.33, p =.002 η^2^ =.81). In contrast, the P2-N1 amplitude difference was significantly larger for older adults for the envelope (t_64_ = 3.29, p =.005, η^2^ =.72), and the beat (t_64_ = 2.11, p = 2.44·10^-5^, η^2^ =1.12). Unlike the other three features, the age-related enhancement of the N1-P1 and N1-P2 for the beat feature (Figure 3E and F) shows later latencies. These findings are consistent with the age-related enhancement of the neural responses to the stimulus onset in Figure 2.

### Sensitivity of neural tracking to musical tempo differs between age groups for early neural responses

Figure 4 shows the TRF time courses (weights) for each of the 13 different musical tempi. The sensitivity of TRF weights to musical tempo was assessed using the slope from the regression of TRF weight as a function of tempo steps at each time point of the time course (Figure 5).

**Figure 4.**
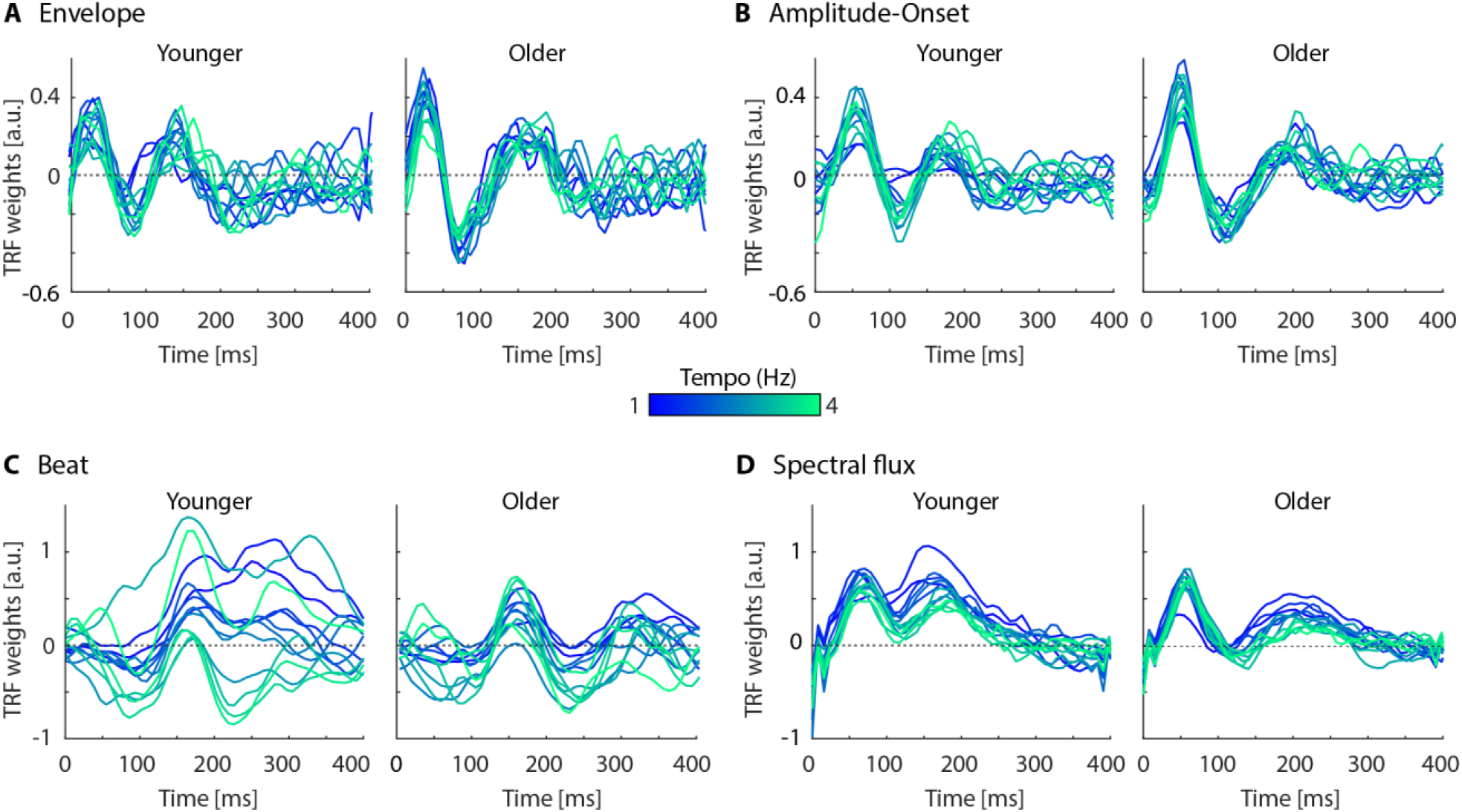
TRF time courses (weights) for each feature in music and music tempo. Each curve depicts the TRF weights from one musical tempo (13 tempi from 1 Hz to 4 Hz in steps of 0.25 Hz, blue to green).

**Figure 5.**
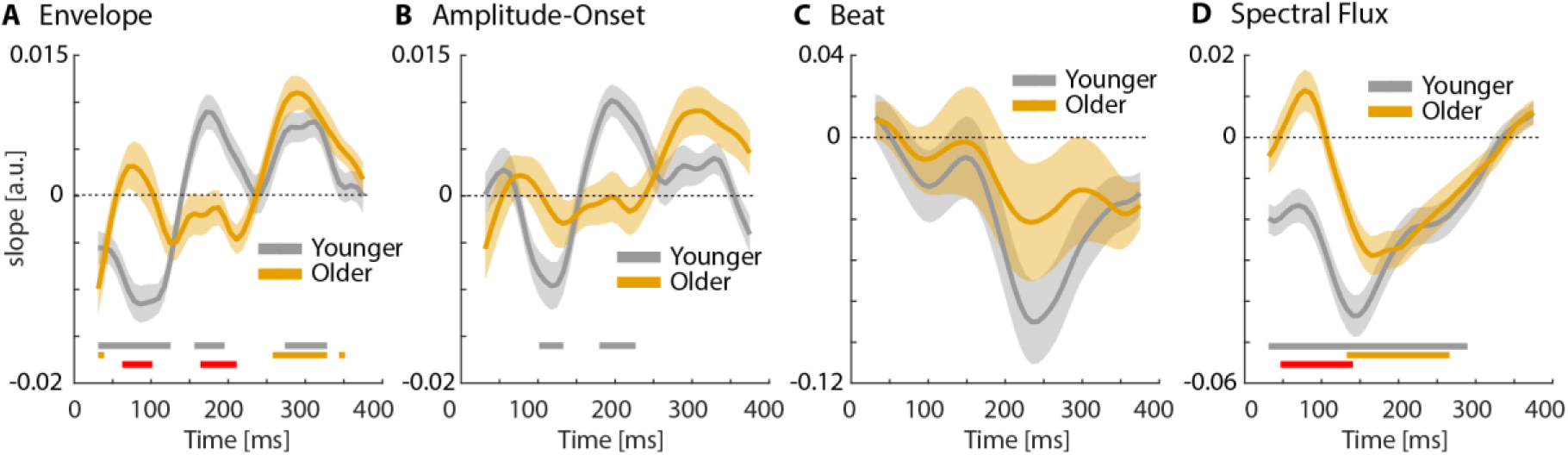
Slopes (linear coefficient) from the regression analysis relating musical tempo to TRF weights. Average slope time courses (shaded area: ±SEM). Horizontal lines at the bottom mark the time window exhibiting a significant difference (pFDR <.05) in the slope between age groups (red line) and within each group against 0 (younger adults: gray; older adults: yellow).

Within each age group, we first examined the TRF sensitivity by comparing slope values against zero, separately for each feature (Figure 5). For this analysis, we were only interested in whether the slopes are different from zero, and were less concerned with the directionality of the effect because the positive and negative deflections in the TRF time courses can influence the slope’s sign while reflecting a similar neural sensitivity to tempo. We evaluate the drivers of any slope effects subsequently in Figure 6 and the corresponding text.

**Figure 6.**
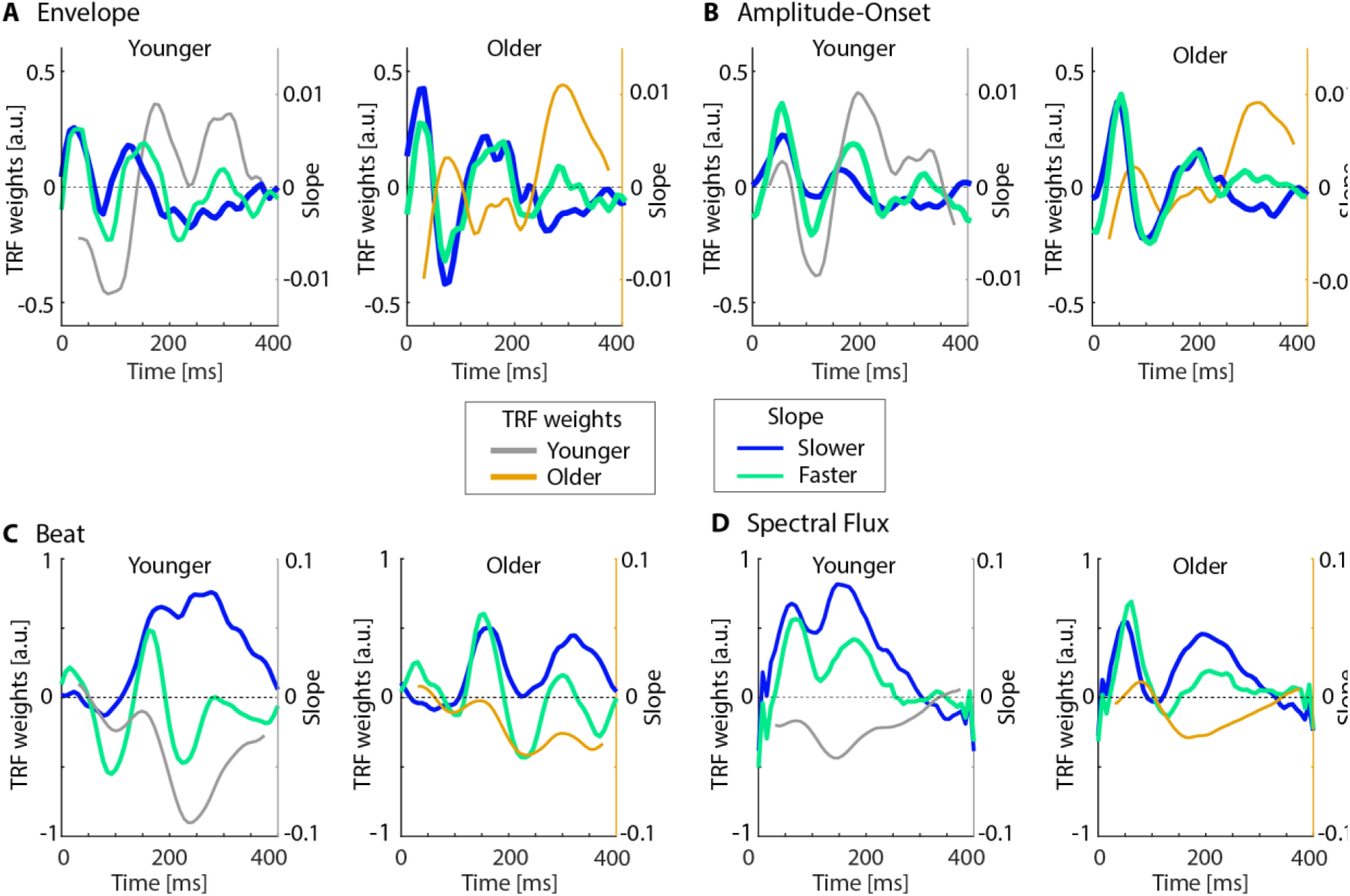
TRF weights for slower and faster tempi, overlaid by the slope (linear coefficient) from the regression model relating TRF weights to musical tempo. A: Average TRF weights across the slowest three tempi (1-1.5Hz; blue lines) and the fastest three tempi (3.5-4Hz; green lines) overlaid by the slope (linear coefficients) from the regression analysis of TRF weights as a function of 13 tempo steps in younger and older adults for the amplitude envelope. B-D: similar to panel A, but for the other three features: amplitude-onset envelope (panels B), beat (panel C) and spectral flux (panel D).

For the amplitude envelope and onset-envelope, younger individuals showed effects of tempo modulation in two early time windows (Figure 5A and B): 100-133 ms (negative values) and 179-226 ms (positive values), but not for older adults. Age-group differences for the amplitude envelope were observed in the 62-102 ms and 164-211 ms ranges (Figure 5A). For spectral flux, younger adults show an extended effect of tempo (32-289 ms), such that TRF weights were larger for slower tempi, whereas older adults showed the effect only for a later time window (133-266 ms). Indeed, the effect was greater in younger compared to older adults in an early time window (47- 140 ms). No effects of tempo nor age group were observed for the beat. In sum, the results from these analyses show that early auditory cortex responses in younger adults are sensitive to musical tempo, whereas only later responses are sensitive to tempo in older adults.

The linear effects characterized by the slopes in Figure 5 reflect neural sensitivity to tempo, but the polarity of the slopes varied due to the TRF deflections in the time courses. To characterize in more detail the underlying structure of the neural tempo sensitivity, we overlaid the averaged TRF time courses across the slowest three tempi (1, 1.25, and 1.5 Hz) and the fastest three tempi (3.5, 3.75, and 4 Hz) with the slope time courses (Figure 6). This indicates that the tempo effect on the envelope and the amplitude-onset in younger adults, at least to some extent, results from a latency delay of the N1 and P2 components as tempo increases. Both age groups showed sensitivity to tempo for the envelope and spectral flux in a later time window, such that the envelope TRF weights were larger for faster tempos, whereas the spectral flux TRF weights were larger for slower tempos.

### Age-groups differences in EEG prediction accuracy manifest strongest for spectral flux

Results for EEG prediction accuracy from both age groups are displayed in Figure 7. The mean prediction accuracy for the four features, independent of musical tempo, did not differ between age groups (main effect of Age Group: F_1,64_ =.09, p =.77, η_p_^2^ =.001; Figure 7A). Averaged EEG prediction accuracy across all tempi was largest for spectral flux, relative to the other acoustic features (effect of Acoustic Feature: F_3,192_ = 185.02, p = 2.17·10^-56^, η_p_^2^ = .74), as indicated by post hoc tests (Bonferroni corrected) comparing prediction accuracy values from spectral flux to all three other features in both age groups (all p < .001). There was no Age Group × Feature interaction (F_3,192_ = 2.14, p =.09, η_p_^2^ =.03).

**Figure 7.**
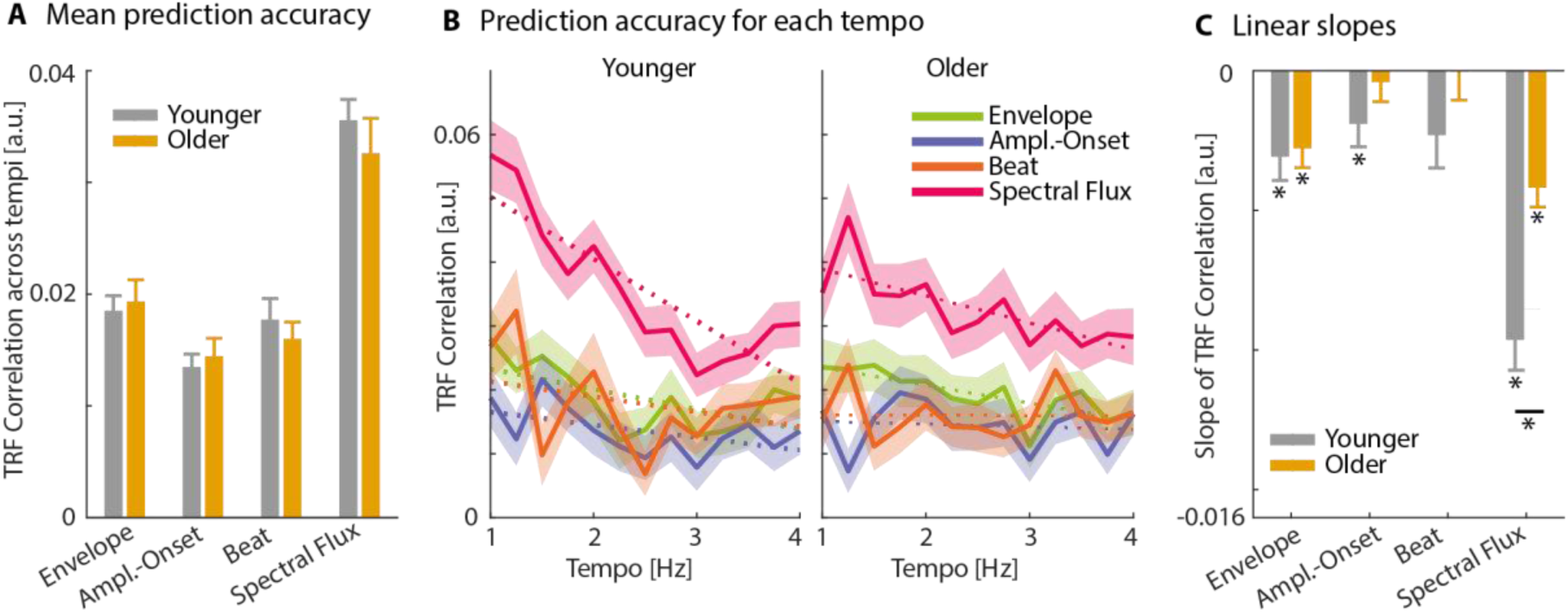
Results for EEG prediction accuracy. A: Mean TRF correlations (± SEM) across 13 tempi in both age groups, separated by four features in naturalistic music. B: Mean TRF (± SEM) correlations plotted against musical tempo separately for four features (represented by four colored lines and shaded areas). Dotted lines denote linear regression of TRF correlations as a function of 13 tempi for each feature. C: Group comparison of the slopes of linear regression for four features in two age groups. The gray and yellow bars represent slope values from younger and older adults, respectively. Asterisks near each bar data denote significant differences of slope values against 0, whereas asterisks above the straight lines denote significant differences between groups. *p<.05.

To explore the potential modulation of EEG prediction accuracy by musical tempo, we fitted a linear function (regression) to prediction accuracies as a function of tempo. Slopes were significantly smaller than zero for the amplitude envelope and spectral flux for both younger and older adults, and for the onset-envelope for younger adults (for all p < .001; Figure 7C), showing that EEG prediction accuracy decreased with increasing musical tempo. The slopes were submitted to an rmANOVA, yielding an effect of Feature (F_3,192_ = 33.64, p = 1.61·10^-17^, η_p_^2^ =.35), Age Group (F_1,64_ = 6.02, p =.02, η_p_^2^=.09), and an interaction between Age Group and Feature (F_3,192_ = 5.56, p =.001, η_p_^2^ =.08). Post hoc Bonferroni-corrected tests revealed a more negative slope for younger compared to older adults for spectral flux (t_64_ = 4.15, p =.002, η^2^ = 1.02), whereas there were no age-group differences for the other features (all p ≥ .10).

## DISCUSSION

In the present study, we examined age-related changes in neural tracking of naturalistic pieces of music that differed in tempo. We focused on features that encapsulate distinct temporal and spectral aspects of naturalistic music (envelope, onset-envelope, beat, and spectral flux). Our findings revealed several key points: spectral flux predicted EEG responses best in both younger and older adults, especially for slower music (1-2 Hz). Critically, older participants showed enhanced neural tracking of all features during early processing stages (0 to 200 ms). However, older adults showed reduced sensitivity to musical tempo compared to younger adults that was apparent for the neural tracking of the envelope, onset-envelope, and spectral flux. The current study suggests complex changes in the neural sensitivity to features in naturalistic music associated with aging.

### Aging is associated with increased onset responses and neural tracking

Older participants exhibited overall larger neural responses to the onset of musical stimuli, particularly reflected in the N1 deflection of the event-related potential (Figure 2). The N1 component peaks between 90 and 130 ms after sound onset and originates from the auditory cortex (Näätänen and Picton, 1987; Scherg et al., 1989; Tremblay et al., 2001; Lyytinen et al., 2002). The observed enhancement of the N1 to sound onset is consistent with previous research showing hyperactivity to sound in older people (Anderer et al., 1996; Amenedo and Díaz, 1999; Herrmann et al., 2013), which is thought to result from increased neural excitability and reduced neural inhibition that follows the reduced sound input associated with peripheral deafferentation (Caspary et al., 2008; Auerbach et al., 2014; Ouellet and de Villers-Sidani, 2014; Zhao et al., 2016; Resnik and Polley, 2017; Herrmann and Butler, 2021). The age-related enhancement is unlikely due to attentional effects as it has been observed for a range of stimuli in the absence of attention in humans (Sörös et al., 2009; Herrmann et al., 2016, 2018) and also in anesthetized animal models (Herrmann et al., 2017; Parthasarathy et al., 2019). The current data may thus suggest that the auditory cortex in older adults exhibited reduced inhibition and heightened excitation levels compared to auditory cortex of younger adults.

Furthermore, the P2 deflection in auditory evoked potentials to music onset was delayed in older compared to younger adults (Figure 2). The P2 component, originating from the auditory association cortex (Wood and Wolpaw, 1982; Whiting et al., 1998; Sharma and Dorman, 1999), is sensitive to complex acoustic features (Bertoli and Probst, 2005; Herrmann et al., 2016). While the exact mechanism for the delayed P2 is unknown, it appears to be a consistent with observations in studies examining the age-related changes in neural responses to sounds (Bertoli et al., 2005; Tremblay and Ross, 2007; Alain et al., 2022).

Our investigation of the TRF time courses revealed that older adults exhibited larger neural tracking for the envelope, amplitude-onset, and spectral flux (Figures 3 and 4), suggesting that hyperactivity also manifests in the neural tracking of relevant features of naturalistic music. This enhancement is also consistent with recent work showing increased neural synchronization of different sound features. For example, neural synchronization with the amplitude envelope and the amplitude-onset envelope of speech is often enhanced in older people compared to younger adults (Presacco et al., 2016; Karunathilake et al. 2023; Panela et al. 2024; McClaskey, 2024). Neural synchronization to low-frequency amplitude modulations of non-speech sounds, such as narrow-band noises, is also enhanced in older people (Purcell et al., 2004; Irsik et al. 2021; Herrmann et al. 2023). Less work exists with regards to the neural synchronization with frequency modulations in sounds, but the evidence that exists also points to enhanced synchronization in older adults (Herrmann et al., 2023), consistent with the results for spectral flux in the current study.

Age-groups did not differ in neural synchronization with the beat (Figure 3C; at least in the FDR-threshold time courses), but beat-based synchronization was the most variable in both age groups (Figure 3C and 4C). The beat may not be the most reliable feature for predicting neural responses in TRF models, possibly because the regular, discrete, and low-frequency nature of the beat occurrence might interfere with the model’s calculations, particularly for the time-lags typically used < 500 ms. Moreover, the perception of the beat is subjective (Danielsen et al., 2024) and the beat identified by the professional percussionist may not necessarily align with the beat perceived by all participants, potentially contributing to the variability. Individual estimates for each participant were not available, but this could be explored further in future TRF work.

Overall, the current study reveals a generalized enhancement of neural responses and neural tracking of features in naturalistic music in older adults, indicating that individual features in music are overly represented in the aged auditory cortex.

### Spectral flux best predicts EEG responses in younger and older adults

Both age groups showed the greatest EEG prediction accuracy (i.e., TRF correlation) for spectral flux, a measure of rapid spectral changes. This prominence likely arises because spectral changes are a key acoustic cue underlying timbre complexity (Alluri et al., 2011; McAdams, 2013) and melody (Zatorre et al., 1994; Janata et al., 2002). Consistent with this, our current and previous work (Weineck et al., 2022) suggest that spectral flux better predicts EEG activity than amplitude features. Processing of spectral flux may dynamically engage tonotopically organized primary cortex (Formisano et al., 2003) and integrative non-primary areas (Norman-Haignere et al., 2015).

### Older adults exhibit reduced sensitivity to tempo in naturalistic music

Both younger and older adults demonstrated greater neural tracking of the amplitude envelope and spectral flux at slower compared to faster tempi, evidenced by higher TRF correlations at 1-2 Hz (associated with the negative slope in Figure 7B and C). This finding aligns with previous research suggesting that slower tempos drive neural synchronization more strongly than higher tempos in music (Doelling and Poeppel, 2015; Weineck et al., 2022), that slow tempos are particularly relevant for the processing of music (Chang et al., 2024), and that slow tempos between 1-2 Hz reflect an optimal information sampling of auditory sequences (Zalta et al., 2020). The results are also consistent with work showing that slow temporal modulations in music are closely linked to the perception of temporal regularities and rhythms, which are essential for music perception (Drake and Botte, 1993; Large and Jones, 1999), and with the frequency at which individuals spontaneously tap (Burger et al., 2018).

Interestingly, older compared to younger adults showed attenuated modulation in EEG prediction accuracy (i.e., TRF correlations) across tempi for spectral flux, and this reduced tempo sensitivity stemmed primarily from reduced prediction accuracy at slower tempi (1-2 Hz; Figure 7B.). TRF temporal waveforms extended this pattern to the amplitude envelope, with age-related reductions in tempo sensitivity localized within early (<200 ms) sensory processing windows (Figures 5 and 6). These early components (N1 and P2), primarily generated in primary auditory cortex (Näätänen and Picton, 1987; Tremblay et al., 2001), may reflect an automatic encoding of acoustic features (Hillyard and Picton, 1978; Picton et al., 1978), and suggest reduced temporal sensitivity in early features extraction stages of the auditory system. The impairment in tempo sensitivity is particularly noteworthy given the older adults’ enhanced responses in this early time window to the same features in naturalistic music (Figure 3). Neural activity from both age groups was similarly sensitive to musical tempo (for envelope and spectral flux) in later time windows (>200 ms), which is perhaps less automatic.

Previous work investigating age-related changes in neural sensitivity to spectral-temporal coherence, amplitude modulations, and frequency modulations in simpler sounds (e.g., narrow-band noises) observed a similar pattern of results for a different neural signature of regularity processing. That is, a low-frequency drift in neural activity – that has been shown to be sensitive to the detection of a regular pattern in sounds (Barascud et al., 2016; Teki et al., 2016; Southwell et al., 2017) – was reduced in older compared to younger adults (Herrmann et al., 2019, 2022, 2023). This reduction in the neural signature of sound regularity processing was present despite the fact that neural responses to sound onsets were enhanced – suggesting hyperactivity – for older adults (Ross and Tremblay, 2009; Bidelman et al., 2014; Irsik et al., 2021). Thus, auditory cortex hyperactivity in aging enhances responses to various features in naturalistic music, but possibly at the cost of reduced sensitivity to acoustic dynamics, such as musical tempo. This neuro-computational tradeoff may limit temporal precision in complex listening environments.

## Conclusions

The current study reveals how aging alters neural tracking of different features of naturalistic music. Older adults exhibited hyperactivity across multiple features, including sound onsets, amplitude envelope, and spectral flux, suggesting a generalized manifestation of a loss of inhibition in the auditory cortex of older people. Crucially, this hyperresponsiveness coexists with diminished neural sensitivity to musical tempo, particularly affecting early sensory processing (<200 ms). Paradoxically, while spectral flux remained the strongest neural predictor in both age groups, older adults showed reduced neural sensitivity of spectral flux to musical tempo (reduced tracking at slower tempi: 1-2 Hz) relative to younger adults. These findings demonstrate that age-related hyperactivity enhances feature encodings in music but may impair neural sensitivity to musical tempo, suggesting complex changes in the aged auditory system.

## AUTHOR CONTRIBUTIONS

**YR**: Methodology, formal analysis, data curation, visualization, writing - original draft, writing - review and editing, project administration. **KW**: Conceptualization, methodology. **MJH**: Conceptualization, methodology, formal analysis, writing – review and editing, project administration, supervision, funding acquisition. **BH**: Conceptualization, methodology, formal analysis, writing – original draft, writing – review and editing, supervision, funding acquisition.

## STATEMENTS AND DECLARATIONS

The authors have no conflicts or competing interests.

## DATA AVAILABILITY

Data will be available to other researchers upon reasonable request to the corresponding author.

## Acknowledgements

The research was supported by the Canada Research Chair program (CRC-2023-00383) and the Natural Sciences and Engineering Research Council of Canada (NSERC Discovery Grant: RGPIN-2021-02602) granted to BH. This work was further supported by a European Research Council Starting Grant (BRAINSYNC) and a Max Planck Research Group granted to MJH. We thank the labs team at the Max Planck for Empirical Aesthetics for technical support and assistance with participant recruitment.

## REFERENCES

1. Alain C, Chow R, Lu J, Rabi R, Sharma VV, Shen D, Anderson ND, Binns M, Hasher L, Yao D, Freedman M (2022) Aging Enhances Neural Activity in Auditory, Visual, and Somatosensory Cortices: The Common Cause Revisited. Journal of Neuroscience 42:264–275.

2. Alain C (2014) Effects of age-related hearing loss and background noise on neuromagnetic activity from auditory cortex. Front Syst Neurosci 8 Available at: 10.3389/fnsys.2014.00008.

3. Alain C, McDonald K, Van Roon P (2012) Effects of age and background noise on processing a mistuned harmonic in an otherwise periodic complex sound. Hear Res 283:126–135.

4. Alluri V, Toiviainen P, Jääskeläinen IP, Glerean E, Sams M, Brattico E (2011) Large-scale brain networks emerge from dynamic processing of musical timbre, key and rhythm. Neuroimage 59:3677–3689.

5. Amenedo E, Díaz F (1999) Ageing-related changes in the processing of attended and unattended standard stimuli. Neuroreport 10:2383–2388.

6. Anari M, Axelsson A, Eliasson A, Magnusson L (1999) Hypersensitivity to sound: Questionnaire data, audiometry and classification. Scand Audiol 28:219–230.

7. Anderer P, Semlitsch HV, Saletu B (1996) Multichannel auditory event-related brain potentials: effects of normal aging on the scalp distribution of N1, P2, N2 and P300 latencies and amplitudes. Electroencephalogr Clin Neurophysiol 99:458–472.

8. Auerbach BD, Rodrigues PV, Salvi RJ (2014) Central Gain Control in Tinnitus and Hyperacusis. Front Neurol 2014 5:206.

9. Barascud N, Pearce MT, Griffiths TD, Friston KJ, Chait M (2016) Brain responses in humans reveal ideal observer-like sensitivity to complex acoustic patterns. Proc. Natl. Acad. Sci. U.S.A. 113:E616–E625.

10. Bauer A-KR, Kreutz G, Herrmann CS (2015) Individual musical tempo preference correlates with EEG beta rhythm. Psychophysiology 52:600–604.

11. Benjamini Y, Hochberg Y (1995) Controlling the False Discovery Rate: A Practical and Powerful Approach to Multiple Testing. J R Stat Soc Series B Stat Methodol 57:289– 300.

12. Bertoli S, Probst R (2005) Lack of standard N2 in elderly participants indicates inhibitory processing deficit. Neuroreport 16:1933–1937.

13. Bertoli S, Smurzynski J, Probst R (2005) Effects of age, age-related hearing loss, and contralateral cafeteria noise on the discrimination of small frequency changes: psychoacoustic and electrophysiological measures. J Assoc Res Otolaryngol 6:207– 222.

14. Bidelman GM, Villafuerte JW, Moreno S, Alain C (2014) Age-related changes in the subcortical-cortical encoding and categorical perception of speech. Neurobiol Aging 35:2526–2540.

15. Bones O, Plack CJ (2015) Losing the music: aging affects the perception and subcortical neural representation of musical harmony. J Neurosci 35:4071–4080.

16. Brodbeck C, Presacco A, Anderson S, Simon JZ (2018) Over-representation of speech in older adults originates from early response in higher order auditory cortex. Acta Acust United Acust 104:774–777.

17. Broderick MP, Di Liberto GM, Anderson AJ, Rofes A, Lalor EC (2021) Dissociable electrophysiological measures of natural language processing reveal differences in speech comprehension strategy in healthy ageing. Sci Rep 11:4963.

18. Burger B, London J, Thompson MR, Toiviainen P (2018) Synchronization to metrical levels in music depends on low-frequency spectral components and tempo. Psychol Res 82:1195–1211.

19. Caspary DM, Ling L, Turner JG, Hughes LF (2008) Inhibitory neurotransmission, plasticity and aging in the mammalian central auditory system. J Exp Biol 211:1781–1791.

20. Chang A, Teng X, Assaneo MF, Poeppel D (2024) The human auditory system uses amplitude modulation to distinguish music from speech. PLoS Biol 22:e3002631.

21. Clinard CG, Tremblay KL, Krishnan AR (2010) Aging alters the perception and physiological representation of frequency: evidence from human frequency-following response recordings. Hear Res 264:48–55.

22. Cohrdes C, Wrzus C, Frisch S, Riediger M (2017) Tune yourself in: Valence and arousal preferences in music-listening choices from adolescence to old age. Dev Psychol 53:1777–1794.

23. Crosse MJ, Di Liberto GM, Bednar A, Lalor EC (2016) The Multivariate Temporal Response Function (mTRF) Toolbox: A MATLAB Toolbox for Relating Neural Signals to Continuous Stimuli. Front Hum Neurosci 10.

24. Crosse MJ, Zuk NJ, Di Liberto GM, Nidiffer AR, Molholm S, Lalor EC (2021) Linear Modeling of Neurophysiological Responses to Speech and Other Continuous Stimuli: Methodological Considerations for Applied Research. Front Neurosci 15:705621.

25. Cruickshanks KJ, Wiley TL, Tweed TS, Klein BE, Klein R, Mares-Perlman JA, Nondahl DM (1998) Prevalence of hearing loss in older adults in Beaver Dam, Wisconsin. The Epidemiology of Hearing Loss Study. Am J Epidemiol 148:879–886.

26. Danielsen A, Brøvig R, Bøhler KK, Câmara GS, Haugen MR, Jacobsen E, Johansson MS, Lartillot O, Nymoen K, Oddekalv KA, Sandvik B, Sioros G, London J (2024) There’s more to timing than time: Investigating musical microrhythm across disciplines and cultures. Music Percept 41:176–198.

27. Decruy L, Vanthornhout J, Francart T (2019) Evidence for enhanced neural tracking of the speech envelope underlying age-related speech-in-noise difficulties. J Neurophysiol 122:601–615.

28. Decruy L, Vanthornhout J, Francart T (2020) Hearing impairment is associated with enhanced neural tracking of the speech envelope. Hear Res 393:107961.

29. Di Liberto GM, Pelofi C, Bianco R, Patel P, Mehta AD, Herrero JL, de Cheveigné A, Shamma S, Mesgarani N (2020) Cortical encoding of melodic expectations in human temporal cortex. Elife 9:e51784.

30. Ding N, Simon JZ (2012) Neural coding of continuous speech in auditory cortex during monaural and dichotic listening. J Neurophysiol 107:78–89.

31. Doelling KB, Poeppel D (2015) Cortical entrainment to music and its modulation by expertise. Proc. Natl. Acad. Sci. U.S.A. 112:E6233–E6242.

32. Drake C, Botte MC (1993) Tempo sensitivity in auditory sequences: evidence for a multiple- look model. Percept Psychophys 54:277–286.

33. Feder K, Michaud D, Ramage-Morin P, McNamee J, Beauregard Y (2015) Prevalence of hearing loss among Canadians aged 20 to 79: Audiometric results from the 2012/2013 Canadian Health Measures Survey. Health Rep 26:18–25.

34. Fiedler L, Wöstmann M, Graversen C, Brandmeyer A, Lunner T, Obleser J (2017) Single- channel in-ear-EEG detects the focus of auditory attention to concurrent tone streams and mixed speech. J Neural Eng 14:036020.

35. Formisano E, Kim DS, Di Salle F, van de Moortele PF, Ugurbil K, Goebel R (2003) Mirror- symmetric tonotopic maps in human primary auditory cortex. Neuron 40:859–869.

36. Friedman D (2013) The cognitive aging of episodic memory: a view based on the event- related brain potential. Front Behav Neurosci 7:111.

37. Friedman D, Simpson G, Hamberger M (1993) Age-related changes in scalp topography to novel and target stimuli. Psychophysiology 30:383–396.

38. Genovese CR, Lazar NA, Nichols T (2002) Thresholding of statistical maps in functional neuroimaging using the false discovery rate. Neuroimage 15:870–878.

39. Goman AM, Lin FR (2016) Prevalence of Hearing Loss by Severity in the United States. Am J Public Health 106:1820–1822.

40. Goossens T, Vercammen C, Wouters J, van Wieringen A (2016) Aging Affects Neural Synchronization to Speech-Related Acoustic Modulations. Front Aging Neurosci 8:133.

41. Goossens T, Vercammen C, Wouters J, van Wieringen A (2019) The association between hearing impairment and neural envelope encoding at different ages. Neurobiol Aging 74:202–212.

42. Gordon-Salant S, Fitzgibbons PJ (1999) Profile of auditory temporal processing in older listeners. J Speech Lang Hear Res 42:300–311.

43. Gratton MA, Vázquez AE (2003) Age-related hearing loss: current research. Curr Opin Otolaryngol Head Neck Surg 11:367–371.

44. Haufe S, Meinecke F, Görgen K, Dähne S, Haynes J-D, Blankertz B, Bießmann F (2014) On the interpretation of weight vectors of linear models in multivariate neuroimaging. Neuroimage 87:96–110.

45. Henry MJ, Herrmann B, Kunke D, Obleser J (2017) Aging affects the balance of neural entrainment and top-down neural modulation in the listening brain. Nat Commun 8:15801.

46. Herrmann B (2025) Minimal background noise enhances neural speech tracking: Evidence of stochastic resonance. Available at: 10.7554/eLife.100830.2.

47. Herrmann B, Buckland C, Johnsrude IS (2019) Neural signatures of temporal regularity processing in sounds differ between younger and older adults. Neurobiol Aging 83:73– 85.

48. Herrmann B, Butler BE (2021) Hearing loss and brain plasticity: the hyperactivity phenomenon. Brain Struct Funct 226:2019–2039.

49. Herrmann B, Henry MJ, Johnsrude IS, Obleser J (2016) Altered temporal dynamics of neural adaptation in the aging human auditory cortex. Neurobiol Aging 45:10–22.

50. Herrmann B, Maess B, Johnsrude IS (2018) Aging Affects Adaptation to Sound-Level Statistics in Human Auditory Cortex. J Neurosci 38:1989–1999.

51. Herrmann B, Maess B, Johnsrude IS (2022) A neural signature of regularity in sound is reduced in older adults. Neurobiol Aging 109:1–10.

52. Herrmann B, Maess B, Johnsrude IS (2023) Sustained responses and neural synchronization to amplitude and frequency modulation in sound change with age. Hear Res 428:108677.

53. Herrmann B, Parthasarathy A, Bartlett EL (2017) Ageing affects dual encoding of periodicity and envelope shape in rat inferior colliculus neurons. Eur J Neurosci 45:299–311.

54. Hillyard SA, Picton TW (1978) On and off components in the auditory evoked potential. Percept Psychophys 24:391–398.

55. Ibrahim BA, Llano DA (2019) Aging and Central Auditory Disinhibition: Is It a Reflection of Homeostatic Downregulation or Metabolic Vulnerability? Brain Sciences 9120351.

56. Irsik VC, Almanaseer A, Johnsrude IS, Herrmann B (2021) Cortical Responses to the Amplitude Envelopes of Sounds Change with Age. J Neurosci 41:5045.

57. Janata P, Birk JL, Van Horn JD, Leman M, Tillmann B, Bharucha JJ (2002) The cortical topography of tonal structures underlying Western music. Science 298:2167–2170.

58. Janata P (2015) Neural basis of music perception. Handb Clin Neurol 129:187–205.

59. Jayakody DMP, Friedland PL, Martins RN, Sohrabi HR (2018) Impact of Aging on the Auditory System and Related Cognitive Functions: A Narrative Review. Front Neurosci 12:125.

60. Kaneshiro B, Nguyen DT, Norcia AM, Dmochowski JP, Berger J (2020) Natural music evokes correlated EEG responses reflecting temporal structure and beat. Neuroimage 214:116559.

61. Karunathilake IMD, Dunlap JL, Perera J, Presacco A, Decruy L, Anderson S, Kuchinsky SE, Simon JZ (2023) Effects of aging on cortical representations of continuous speech. J Neurophysiol 129:1359–1377.

62. Keitel A, Pelofi C, Guan X, Watson E, Wight L, Allen S, Mencke I, Keitel C, Rimmele J (2025) Cortical and behavioral tracking of rhythm in music: Effects of pitch predictability, enjoyment, and expertise. Ann N Y Acad Sci 1546:120–135.

63. Koelsch S, Gunter T, Friederici AD, Schröger E (2000) Brain Indices of Music Processing: “Nonmusicians” are Musical. J Cogn Neurosci. 12:520–541.

64. Koelsch S (2011) Toward a neural basis of music perception - a review and updated model. Front Psychol 2:110.

65. Lalor EC, Foxe JJ (2009) Neural responses to uninterrupted natural speech can be extracted with precise temporal resolution. Eur J Neurosci 31:189–193.

66. Lalor EC, Power AJ, Reilly RB, Foxe JJ (2009) Resolving precise temporal processing properties of the auditory system using continuous stimuli. J Neurophysiol 102:349–359.

67. Large EW, Jones MR (1999) The dynamics of attending: How people track time-varying events. Psychological Review 106:119–159.

68. Lopez-Poveda EA, Meddis R (2001) A human nonlinear cochlear filterbank. J Acoust Soc Am 110:3107–3118.

69. Luck SJ (2014) An Introduction to the Event-Related Potential Technique, second edition. MIT Press.

70. Luo H, Poeppel D (2007) Phase patterns of neuronal responses reliably discriminate speech in human auditory cortex. Neuron 54:1001–1010.

71. Lyytinen H, Naatanen R, Sokolov EN, Spinks J (2002) The Orienting Response in Information Processing. Psychology Press.

72. Martin BA, Tremblay KL, Korczak P (2008) Speech evoked potentials: from the laboratory to the clinic. Ear Hear 29:285–313.

73. McAdams S (2013) Musical timbre perception. The psychology of music, 3rd ed:35–67 Available at: 10.1016/B978-0-12-381460-9.00002-X.

74. McClaskey CM (2024) Neural hyperactivity and altered envelope encoding in the central auditory system: Changes with advanced age and hearing loss. Hearing Research 442:1–13 Available at: 10.1016/j.heares.2023.108945.

75. Näätänen R, Paavilainen P, Alho K, Reinikainen K, Sams M (1987) The mismatch negativity to intensity changes in an auditory stimulus sequence. Electroencephalogr Clin Neurophysiol Suppl 40:125–131.

76. Näätänen R, Paavilainen P, Rinne T, Alho K (2007) The mismatch negativity (MMN) in basic research of central auditory processing: a review. Clin Neurophysiol 118:2544–2590.

77. Näätänen R, Picton T (1987) The N1 wave of the human electric and magnetic response to sound: a review and an analysis of the component structure. Psychophysiology 24:375– 425.

78. Norman-Haignere S, Kanwisher NG, McDermott JH (2015) Distinct Cortical Pathways for Music and Speech Revealed by Hypothesis-Free Voxel Decomposition. Neuron 88:1281–1296.

79. Ouellet L, de Villers-Sidani E (2014) Trajectory of the main GABAergic interneuron populations from early development to old age in the rat primary auditory cortex. Front Neuroanat 2014 Jun 2;8:40.

80. Panela RA, Copelli F, Herrmann B (2024) Reliability and generalizability of neural speech tracking in younger and older adults. Neurobiol Aging 134:165–180.

81. Parry LV, Maslin MRD, Schaette R, Moore DR, Munro KJ (2018) Increased auditory cortex neural response amplitude in adults with chronic unilateral conductive hearing impairment. Hear Res 372:10–16.

82. Parthasarathy A, Herrmann B, Bartlett EL (2019) Aging alters envelope representations of speech-like sounds in the inferior colliculus. Neurobiol Aging 73:30–40.

83. Patterson RD, Nimmo-Smith I, Holdsworth J, Rice P, Others (1987) An efficient auditory filterbank based on the gammatone function. In: a meeting of the IOC Speech Group on Auditory Modelling at RSRE.

84. Peelle JE, Troiani V, Grossman M, Wingfield A (2011) Hearing loss in older adults affects neural systems supporting speech comprehension. J Neurosci 31:12638–12643.

85. Picton T (2013) Hearing in time: evoked potential studies of temporal processing. Ear Hear 34:385–401.

86. Picton TW, John MS, Dimitrijevic A, Purcell D (2003) Human auditory steady-state responses. Int J Audiol 42:177–219.

87. Picton TW, Woods DL, Proulx GB (1978) Human auditory sustained potentials. I. The nature of the response. Electroencephalogr Clin Neurophysiol 45:186–197.

88. Presacco A, Simon JZ, Anderson S (2016) Evidence of degraded representation of speech in noise, in the aging midbrain and cortex. J Neurophysiol 116:2346–2355.

89. Purcell DW, John SM, Schneider BA, Picton TW (2004) Human temporal auditory acuity as assessed by envelope following responses. J Acoust Soc Am 116:3581–3593.

90. Resnik J, Polley DB (2017) Fast-spiking GABA circuit dynamics in the auditory cortex predict recovery of sensory processing following peripheral nerve damage. Elife 6:e21452.

91. Ross B, Tremblay K (2009) Stimulus experience modifies auditory neuromagnetic responses in young and older listeners. Hear Res 248:48–59.

92. Salvi R, Sun W, Ding D, Chen G-D, Lobarinas E, Wang J, Radziwon K, Auerbach BD (2016) Inner Hair Cell Loss Disrupts Hearing and Cochlear Function Leading to Sensory Deprivation and Enhanced Central Auditory Gain. Front Neurosci 10:621.

93. Scherg M, Vajsar J, Picton TW (1989) A source analysis of the late human auditory evoked potentials. J Cogn Neurosci 1:336–355.

94. Shahin A, Bosnyak DJ, Trainor LJ, Roberts LE (2003) Enhancement of Neuroplastic P2 and N1c Auditory Evoked Potentials in Musicians. J Neurosci 23:5545–5552.

95. Sharma A, Dorman MF (1999) Cortical auditory evoked potential correlates of categorical perception of voice-onset time. J Acoust Soc Am 106:1078–1083.

96. Shehabi AM, Prendergast G, Guest H, Plack CJ (2022) The Effect of Lifetime Noise Exposure and Aging on Speech-Perception-in-Noise Ability and Self-Reported Hearing Symptoms: An Online Study. Front Aging Neurosci 14:890010.

97. Slade K, Plack CJ, Nuttall HE (2020) The Effects of Age-Related Hearing Loss on the Brain and Cognitive Function. Trends Neurosci 43:810–821.

98. Southwell R, Baumann A, Gal C, Barascud N, Friston K, Chait M (2017) Is predictability salient? A study of attentional capture by auditory patterns. Philos Trans R Soc Lond B Biol Sci 2017 Feb 19;372(1714):20160105.

99. Sörös P, Teismann IK, Manemann E, Lütkenhöner B (2009) Auditory temporal processing in healthy aging: a magnetoencephalographic study. BMC Neurosci 10:34.

100. Stewart L, von Kriegstein K, Warren JD, Griffiths TD (2006) Music and the brain: disorders of musical listening. Brain 129:2533–2553.

101. Sun W, Deng A, Jayaram A, Gibson B (2012) Noise exposure enhances auditory cortex responses related to hyperacusis behavior. Brain Res 1485:108–116.

102. Teki S, Barascud N, Picard S, Payne C, Griffiths TD, Chait M (2016) Neural Correlates of Auditory Figure-Ground Segregation Based on Temporal Coherence. Cereb Cortex 26:3669–3680.

103. Tervaniemi M, Rytkönen M, Schröger E, Ilmoniemi RJ, Näätänen R (2001) Superior formation of cortical memory traces for melodic patterns in musicians. Learn Mem 8:295–300.

104. Tremblay K, Kraus N, McGee T, Ponton C, Otis B (2001) Central auditory plasticity: changes in the N1-P2 complex after speech-sound training. Ear Hear 22:79–90.

105. Tremblay K, Ross B (2007) Effects of age and age-related hearing loss on the brain. J Commun Disord 40:305–312.

106. Wagner M, Shafer VL, Haxhari E, Kiprovski K, Behrmann K, Griffiths T (2017) Stability of the Cortical Sensory Waveforms, the P1-N1-P2 Complex and T-Complex, of Auditory Evoked Potentials. J Speech Lang Hear Res 60:2105–2115.

107. Weineck K, Wen OX, Henry MJ (2022) Neural synchronization is strongest to the spectral flux of slow music and depends on familiarity and beat salience. Elife 11:e75515.

108. Whiting KA, Martin BA, Stapells DR (1998) The effects of broadband noise masking on cortical event-related potentials to speech sounds /ba/ and /da/. Ear Hear 19:218–231.

109. Wood CC, Wolpaw JR (1982) Scalp distribution of human auditory evoked potentials. II. Evidence for overlapping sources and involvement of auditory cortex. Electroencephalogr Clin Neurophysiol 54:25–38.

110. Woods TM, Lopez SE, Long JH, Rahman JE, Recanzone GH (2006) Effects of stimulus azimuth and intensity on the single-neuron activity in the auditory cortex of the alert macaque monkey. J Neurophysiol 96:3323–3337.

111. Wöstmann M, Herrmann B, Wilsch A, Obleser J (2015) Neural alpha dynamics in younger and older listeners reflect acoustic challenges and predictive benefits. J Neurosci 35:1458–1467.

112. Zalta A, Petkoski S, Morillon B (2020) Natural rhythms of periodic temporal attention. Nat Commun 11:1051.

113. Zatorre RJ, Evans AC, Meyer E (1994) Neural mechanisms underlying melodic perception and memory for pitch. J Neurosci 14:1908–1919.

114. Zhao Y, Song Q, Li X, Li C (2016) Neural Hyperactivity of the Central Auditory System in Response to Peripheral Damage. Neural Plast 2016:2162105.

115. Zuk NJ, Murphy JW, Reilly RB, Lalor EC (2021) Envelope reconstruction of speech and music highlights stronger tracking of speech at low frequencies. PLoS Comput Biol 17:e1009358.

